# Regulator of Gene Silencing-Calmodulin associates with mRNA granules and the autophagy protein ATG8

**DOI:** 10.1101/858092

**Authors:** W. Craig Conner, Ansul Lokdarshi, Daniel M. Roberts

## Abstract

**Abstract**

Regulator-of-gene-silencing calmodulins (rgsCaM) represent a phylogenetic subfamily of calmodulin-like calcium sensors that are targets of viral induced suppression of posttranscriptional gene silencing by secondary siRNAs. The present work shows that a stress (hypoxia) that induces mRNP granule formation also induces the relocalization of rgsCaM to cytosolic granule-like foci that interact with the surface of stress granule and processing body structures. Co-expression of rgsCaM and its binding protein Suppressor of Gene Silencing 3 causes re-localization and integration of rgsCaM into stress granule structures. RgsCaMs contain a conserved topology that consists for four EF hand like domains (three functional and one divergent) that are separated into two calcium binding lobes with an extended amino terminal region. RgsCaM also contains an “ATG8 family interacting motif” (AIM) within its amino-terminal domain that is characteristic of selective autophagy cargo receptors. Co-localization experiments and ratiometric BiFC analyses in *Nicotiana benthamiana* support the hypothesis that rgsCaM binds directly to ATG8e through this conserved AIM domain, and the two proteins co-localize with mRNP granule markers. Previous reports show that rgsCaM mediates the suppression of gene silencing, at least in part, via turnover of SGS3 via autophagy. A model is proposed for rgsCaM-like proteins as potential mediators of selective autophagy of RNA granules in response to biotic and abiotic stresses.

## Introduction

“Regulator of Gene Silencing Calmodulin” (RgsCaM) is a calmodulin-like EF hand protein that was originally identified in *Nicotiana tabcum* as a binding target of the potyviral RNA silencing (VSR) protein “helper component proteinase” (HC-Pro), (Anandalakshmi et al., 2000). HC-Pro assists in viral evasion of the host defense response by suppressing post transcriptional gene silencing (PTGS) mediated by small interfering RNA silencing pathways of the plant host (Mallory et al., 2001). Transgenic expression of HC-Pro induces the expression of rgsCaM, and rgsCaM directly binds to HC-Pro by yeast two-hybrid analysis (Anandalakshmi et al., 2000). RgsCaM overexpression on its own in the absence of HC-Pro or viral infection prevents or reverses PTGS (Anandalakshmi et al., 2000). It was proposed that rgsCaM is an endogenous suppressor of PTGS, and that viral HC-Pro may take advantage of this physiological function of rgsCaM to suppress antiviral gene silencing and promote escape from the host defense response (Anandalakshmi et al., 2000).

Since its discovery as a target for potyviral HC-Pro, additional studies show that rgsCaM-like proteins interact with multiple viral suppressor proteins from a wide array of plant viruses (Nakahara et al., 2012; Li et al., 2014; Yong Chung et al., 2014; Li et al., 2017). By using surface plasmon resonance analysis, (Nakahara et al., 2012) demonstrated the interaction of tobacco rgsCaM with the bromoviral suppressor of silencing 2b protein as well as other viral suppressors. Moreover, *N. benthamiana* rgsCaM is induced by the DNA geminivirus suppressor of silencing, βC1, and is essential for conferring the suppressor activity of this viral protein (Li et al., 2014). Moreover, rgsCaM exhibits endogenous suppressor activity on its own upon overexpression in *N. benthamiana* (Li et al., 2014). Similarly, the Arabidopsis rgsCaM homolog, *CML39* is induced by the DNA geminivirus tomato golden mosaic virus (TGMV) suppressor of silencing AL2, and interacts directly with the AL2 protein (Yong Chung et al., 2014). Consistent with a proposed role as a target for viral suppressors of silencing, *CML39* overexpression lines exhibited enhanced TGMV geminivirus infectivity, while reduction of *CML39* expression exhibited reduced infectivity (Yong Chung et al., 2014). Collectively, these studies indicate that rgsCaMs may be a common target for viral proteins in mediating PTGS suppressor activity, as well as serving as endogenous regulators of PTGS.

A growing body of evidence suggests that rgsCaM mediates its biological effects by targeting key RNA silencing proteins for degradation by autophagy (Yang et al., 2019). Autophagy is a cytosolic process in which bulk cytosol, selected cargo, or organelles are engulfed in a double membrane and fused to the vacuole/lysosome for degradation and recycling of critical resources during cellular stress (Stolz et al., 2014; Yang and Bassham, 2015). The first connection of rgsCaM to autophagy was provided by (Nakahara et al., 2012) who showed that autophagic inhibitors and RNA_i_ suppression of *Autophagy-related gene 6 (ATG6)* blocked cellular turnover of rgsCaM protein as well as HC-Pro and other viral suppressors of gene silencing.

A strong connection of rgsCaM to autophagy has been discovered by Li et al. (2017), who demonstrated that the SGS3 protein encoded by *SUPPRESSOR OF GENE SILENCING 3* is a direct interaction target for rgsCaM (Li et al., 2017). SGS3 is an RNA-binding homodimeric protein that forms a stable complex with RNA-dependent RNA polymerase 6 (RDR6). SGS3 is an essential chaperone protein for RDR6-catalyzed synthesis of dsRNA templates required for secondary siRNA production in viral and transgene PTGS pathways (Mourrain et al., 2000; Fukunaga and Doudna, 2009; Li et al., 2017). SGS3 and RDR6 accumulate within cytosolic foci termed siRNA bodies that are proposed to be involved in transacting siRNA biosynthesis (Elmayan et al., 2009; Kumakura et al., 2009; Jouannet et al., 2012). Overexpression of rgsCaM lowered the levels of SGS3 protein as well as the number of siRNA bodies (Li et al., 2017). This effect of rgsCaM overexpression was suppressed by autophagy inhibitors and RNA_i_ reduction of the expression of key autophagy genes (Li et al., 2017). From these observations, the suppression of PTGS activity by rgsCaM was attributed to the autophagic turnover of SGS3/RDR6, which would reduce the ability of the cell to produce dsRNA for the production of secondary siRNAs. In support of this hypothesis, it has been observed that rgsCaM preferentially suppresses secondary siRNA silencing pathways (e.g., sense transgene and viral) and not primary silencing pathways (e.g., inverted repeat hairpin transgenes) that operate independent of an RDR pathway (Nakamura et al., 2014).

The question of endogenous physiological functions of rgsCaM proteins remains open. Potential insight into a role in abiotic stress responses and mRNA ribostasis comes from the observation that the rgsCaM-like protein CML38 from Arabidopsis is induced during flooding and hypoxia stress where it accumulates within cytosolic ribonucleoprotein structures known as stress granules (SGs) (Lokdarshi et al., 2016). SGs represent non-membranous cytosolic granule assemblies consisting of arrested pre-initiation mRNA complexes that accumulate during various cellular stresses (Protter and Parker, 2016; Chantarachot and Bailey-Serres, 2018). In the present study, the association of rgsCaM with hypoxia stress-induced granule structures was analyzed. Further, evidence for direct association of rgsCaM with the autophagic receptor ATG8 through a canonical AIM domain is presented. The significance of these observations and a model for rgsCaM and related proteins as potential mediators of granule turnover by autophagy (granulophagy) under conditions of cellular stress is presented.

## Results

### RgsCaM localizes to hypoxia-induced cytosolic foci

To investigate the subcellular localization of tobacco rgsCaM, *Nicotiana benthamiana* leaves were transiently transfected with an expression construct consisting of a translational fusion of rgsCaM with C-terminal yellow fluorescent protein (YFP) (Fig. 1). RgsCaM accumulates in distinct cytosolic foci as well as within the nuclear compartment (Fig. 1). In previous work, it was shown that submergence of *N. benthamiana* leaf disks in aqueous media induces a hypoxic state that leads to a time-dependent formation of cytosolic SGs (Weber et al. 2008; Sorenson and Bailey-Serres, 2014; Lokdarshi et al., 2016). A time series of confocal microscopy images after tissue submergence (Fig. 2) shows that rgsCaM initially shows a diffuse cytosolic signal and that over time the protein redistributes into cytosolic foci.

**Fig 1.**
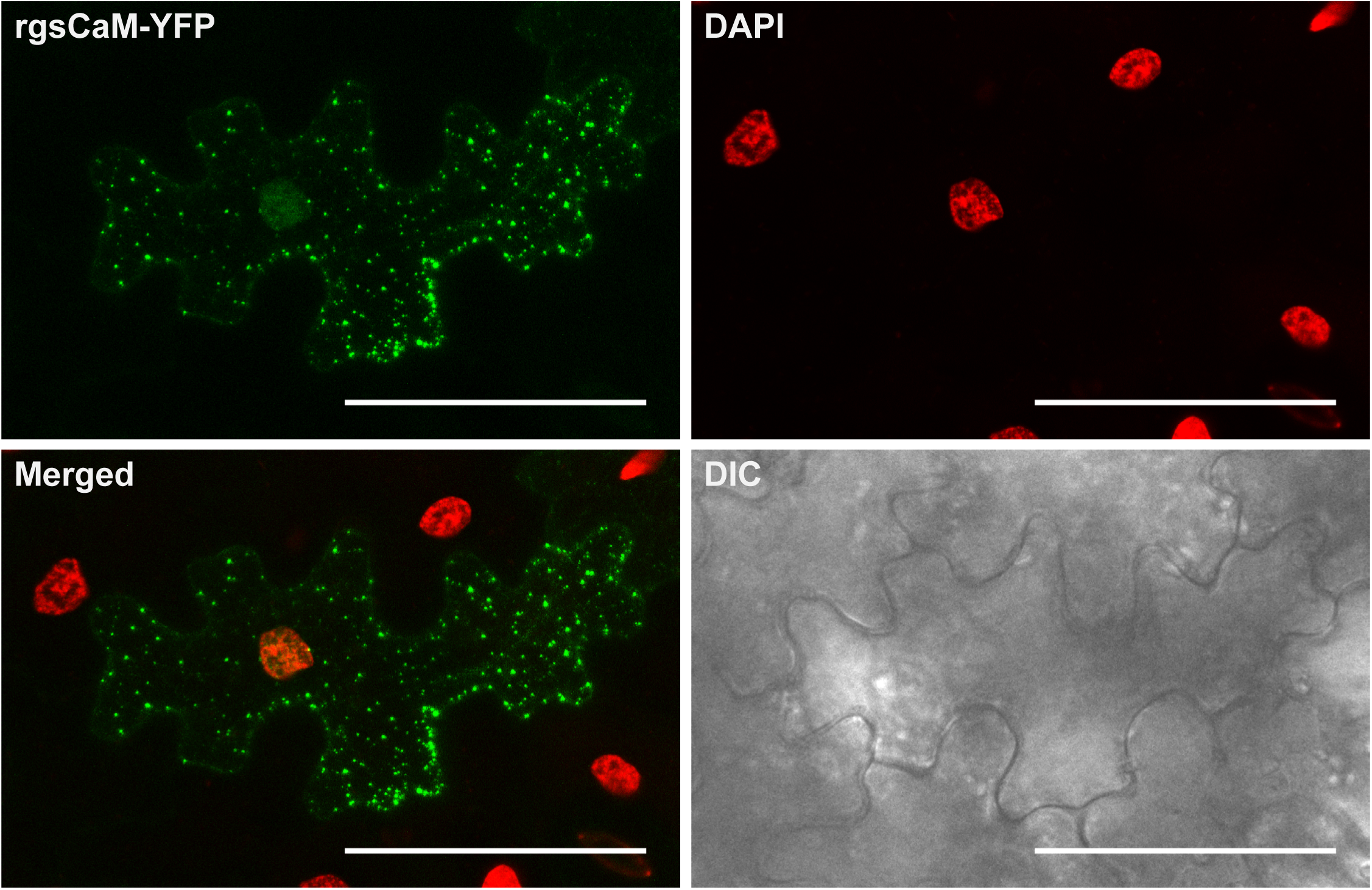
Localization of rgsCaM-YFP in cytosolic granule-like foci in *N benthamiana* leaf cells. *N benthamiana* leaves were transfected with a *rgsCaM-YFP* expression construct by *Agrobacterium* infiltration, and were subjected to submergence-induced hypoxia as described in the Materials and Methods. RgsCaM-YFP was visualized by confocal microscopy, and nuclei were stained with DAPI. Scale bars are 50 µm.

**Fig 2.**
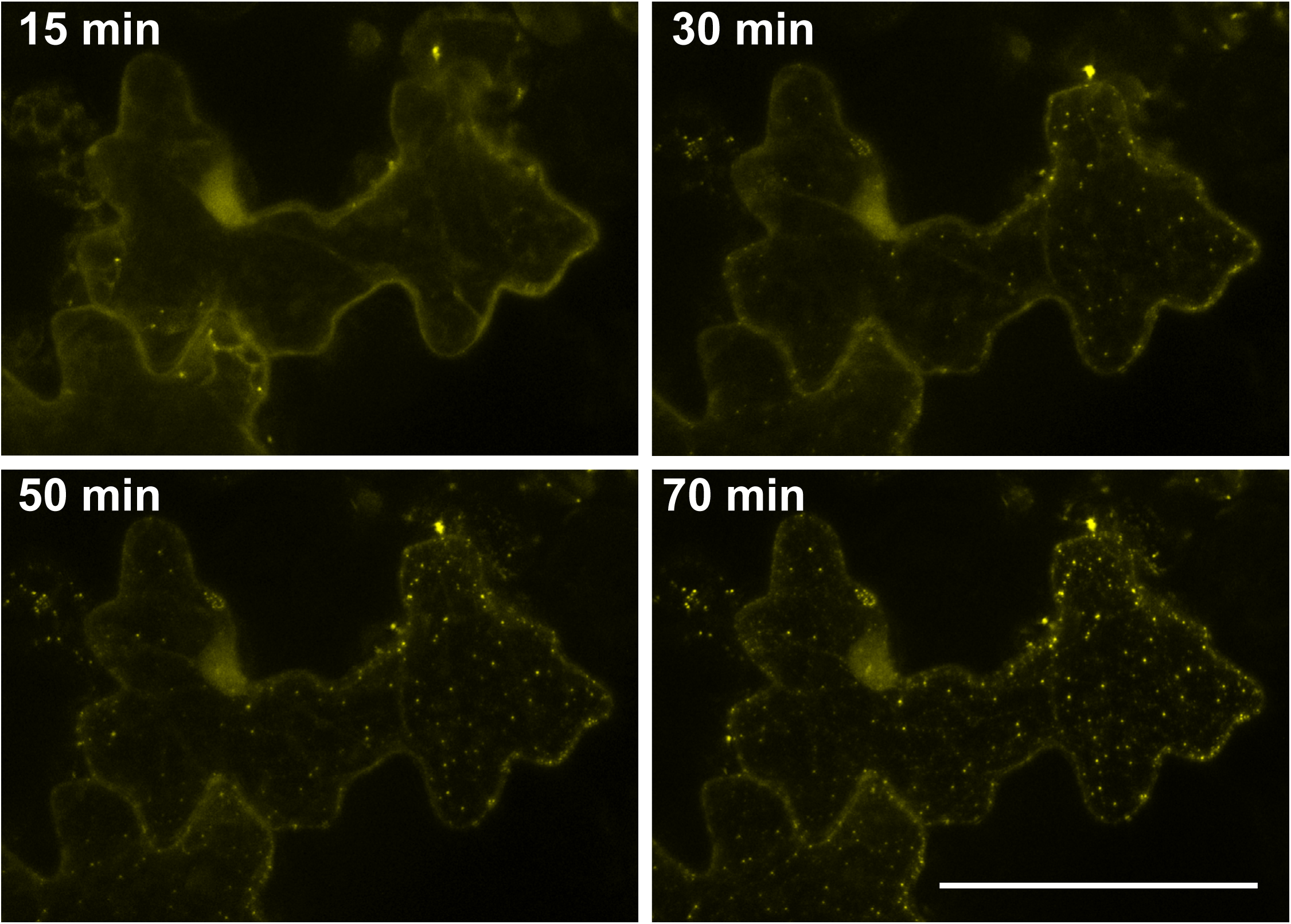
Timecourse of. RgsCaM-YFP cytosolic foci formation in response to submergence-induced hypoxia. A representative cell from a transfected *N. benthamiana* leaf expressing rgsCaM-YFP was imaged over the indicated times after submergence. Scale bar is 50µm.

### RgsCaM granules are independent structures that associate at the periphery of SGs and PBs under hypoxia

Given the previous observation that the rgsCaM-like protein CML38 associates with hypoxia-induced SGs (Weber et al., 2008; Lokdarshi et al., 2016), the co-localization of rgsCaM with the SG marker RBP47B was investigated. *N. benthamiana* leaves were co-transfected with constructs expressing rgsCaM-YFP and RBP47B-CFP and imaging was performed in submerged leaf discs (Fig. 3). RBP47B accumulates within cytosolic foci consistent with the appearance of SGs as well as within the nucleus (Fig. 3A), similar to previous observations (Lokdarshi et al., 2016). RgsCaM cytocolic foci are mostly distinct from the SG marker RBP47B (Fig. 3A). This is in contrast to previous analyses of Arabidopsis CML38 (Lokdarshi et al., 2016), but is supported by co-localization quantification (Manders et al., 1992), that showed low Manders’ co-efficients for RBP47B overlapping with rgsCaM (MCC_RBP47B:rgsCaM_ = 0.12), as well as for rgsCaM overlapping with RBP47B (MCC_rgsCaM:RBP47B_ = 0.18).

**Fig 3.**
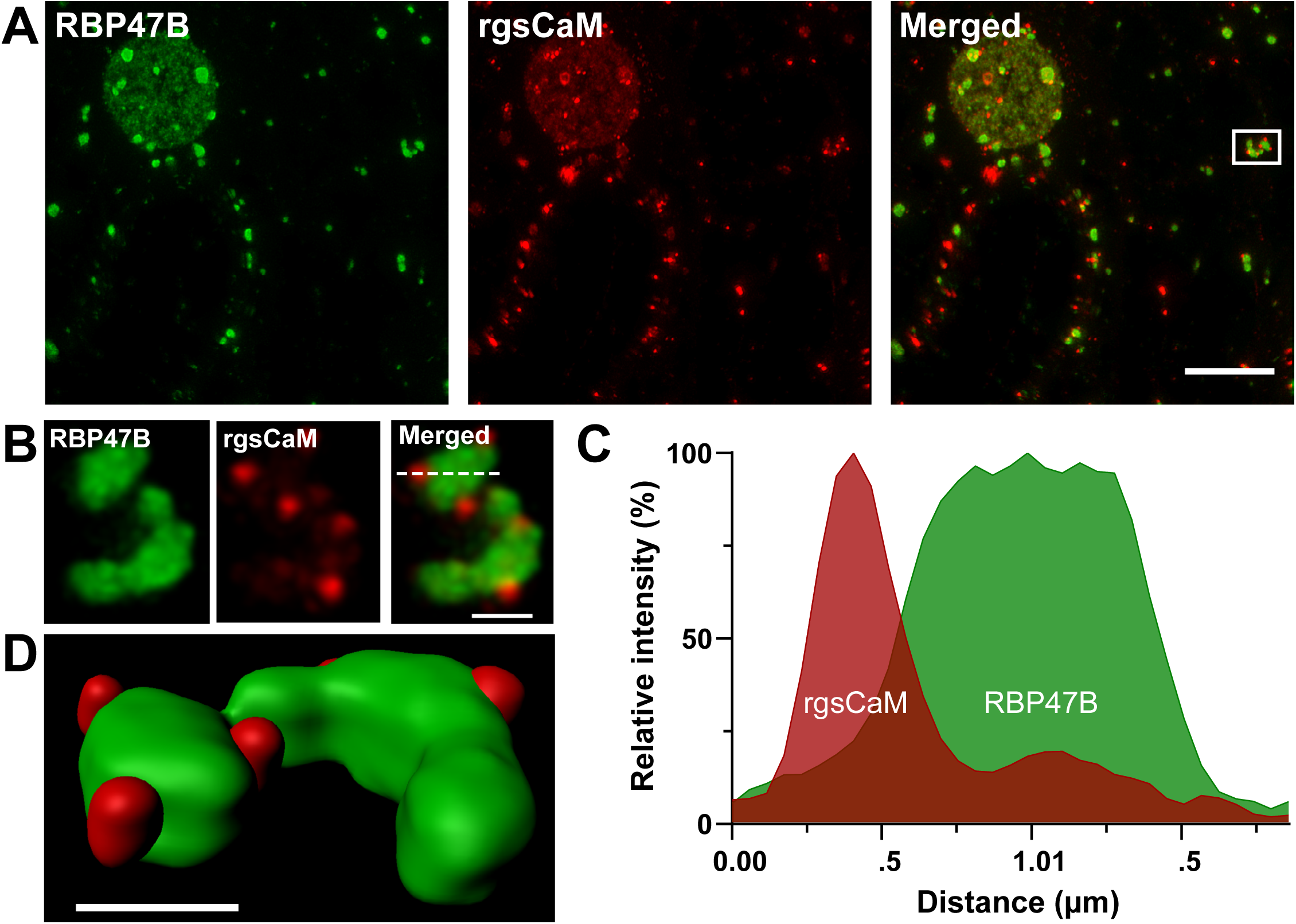
Co-localization of rgsCaM-YFP and stress granule markers in hypoxic *N. benthamiana* leaves. Co-localization experiments with *N. benthamiana* leaves co-transfected with constructs that express rgsCaM-YFP and the stress granule marker RBP47B-CFP. **A)** 2D projections from a representative cell, consisting of 99 optical sections with a step size of 0.3 µm, and subjected to standard deconvolution as described in the Materials and Methods. Scale bars are 10 µm. **B)** Magnification of the boxed region shown in panel A. Scale bar is 1µm. A smoothing filter was applied in Leica LasX software. **C)** A line plot profile of fluorescence intensities calculated along the axis shown by the dashed line in panel B. **D)** 3D model reconstruction from the dataset shown in B using the Imaris Software (see also Fig. S1). Scale bar is 1 µm.

However, a closer examination of the SG structures showed the apparent association of smaller rgsCaM foci at the periphery of SGs (Fig. 3B). A line plot profile (Fig. 3C) shows that while the rgsCaM and RBP47B fluorescence intensity maxima are clearly distinct, in some cases the two particles appear to associate with partially overlapping fluorescent signals. A three-dimensional model was generated from the volumetric signal intensities in the Z-stack using Imaris software (Fig. 3D, Fig. S1). The resulting model suggests that rather than localizing within SGs, rgsCaM appears to interact with the surface of SGs.

A similar analysis was performed with rgsCaM and the processing body (PB) marker decapping protein 1 (DCP1) (Fig. 4). Leaf disc submergence resulted in the typical localization of rgsCaM-YFP to small cytosolic foci, with additional apparent co-localization with larger DCP1-CFP PB foci (Fig. 4). This dual localization of rgsCaM is reflected by Manders co-localization analysis The MCC_DCP1:rgsCaM_ for DCP1 overlapping with rgsCaM is 0.96, while the MCC_rgsCaM:DCP1_ for rgsCaM overlapping with DCP1 is 0.08. This observation suggests that most PBs are associated with rgsCaM bodies, but the majority of rgsCaM bodies are independent from DCP1-containing processing bodies. A closer examination at high resolution with deconvolution reveals substructure to the rgsCaM/DCP1 aggregates, and distinct localization of rgsCaM bodies and DCP1 stained PB (Fig. 5). Three-dimensional modeling using Imaris through a representative PB show that rgsCaM bodies appear to a form a ring that surrounds the PB (Fig. 5D, Fig. S2).

**Fig 4.**
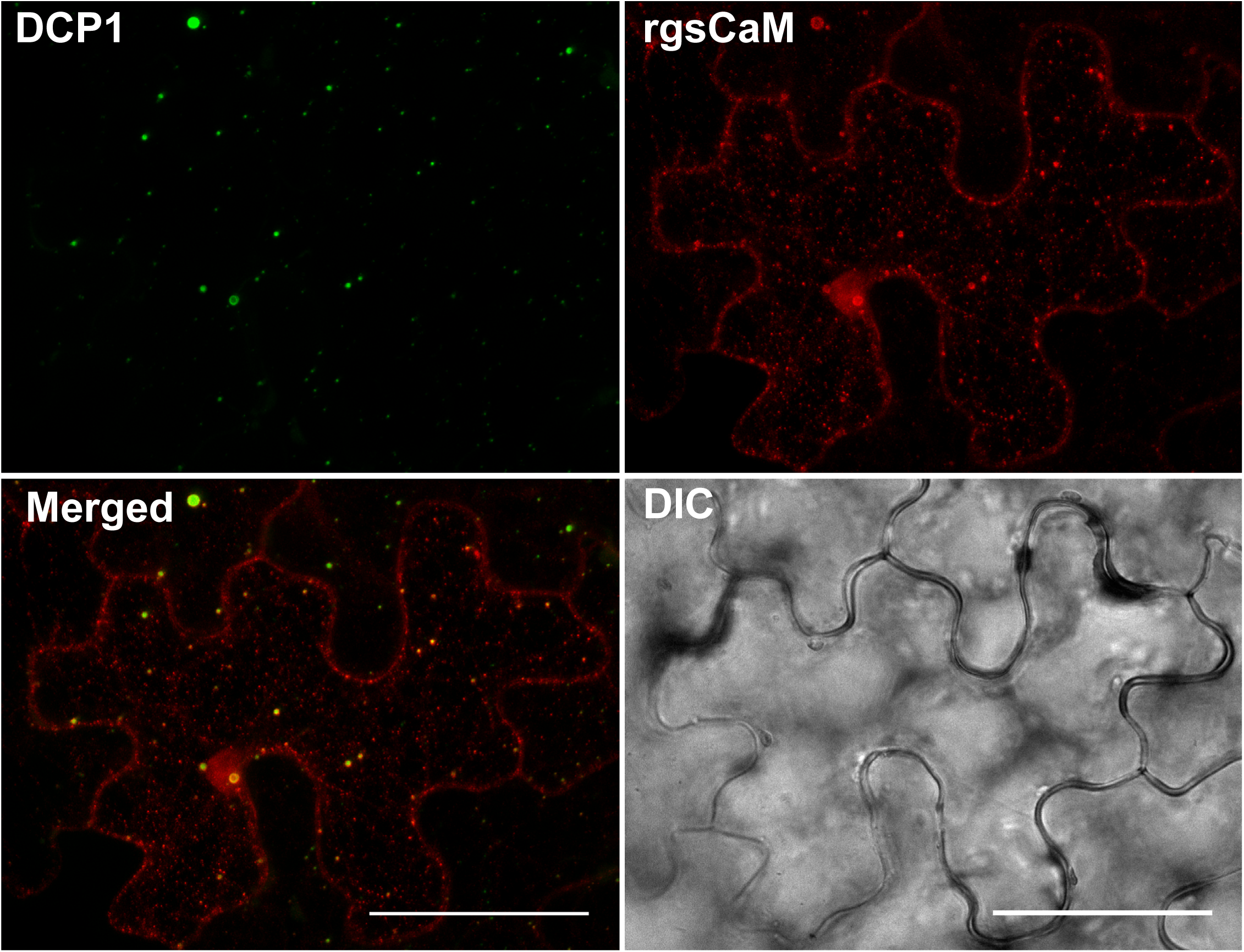
Co-localization of rgsCaM-YFP and processing body markers in hypoxic *N. benthamiana* leaves. Leaf sections of *N. benthamiana* leaves co-transfected with constructs that express rgsCaM-YFP and the processing body marker DCP1-CFP. Scale bar is 50 µm.

**Fig 5.**
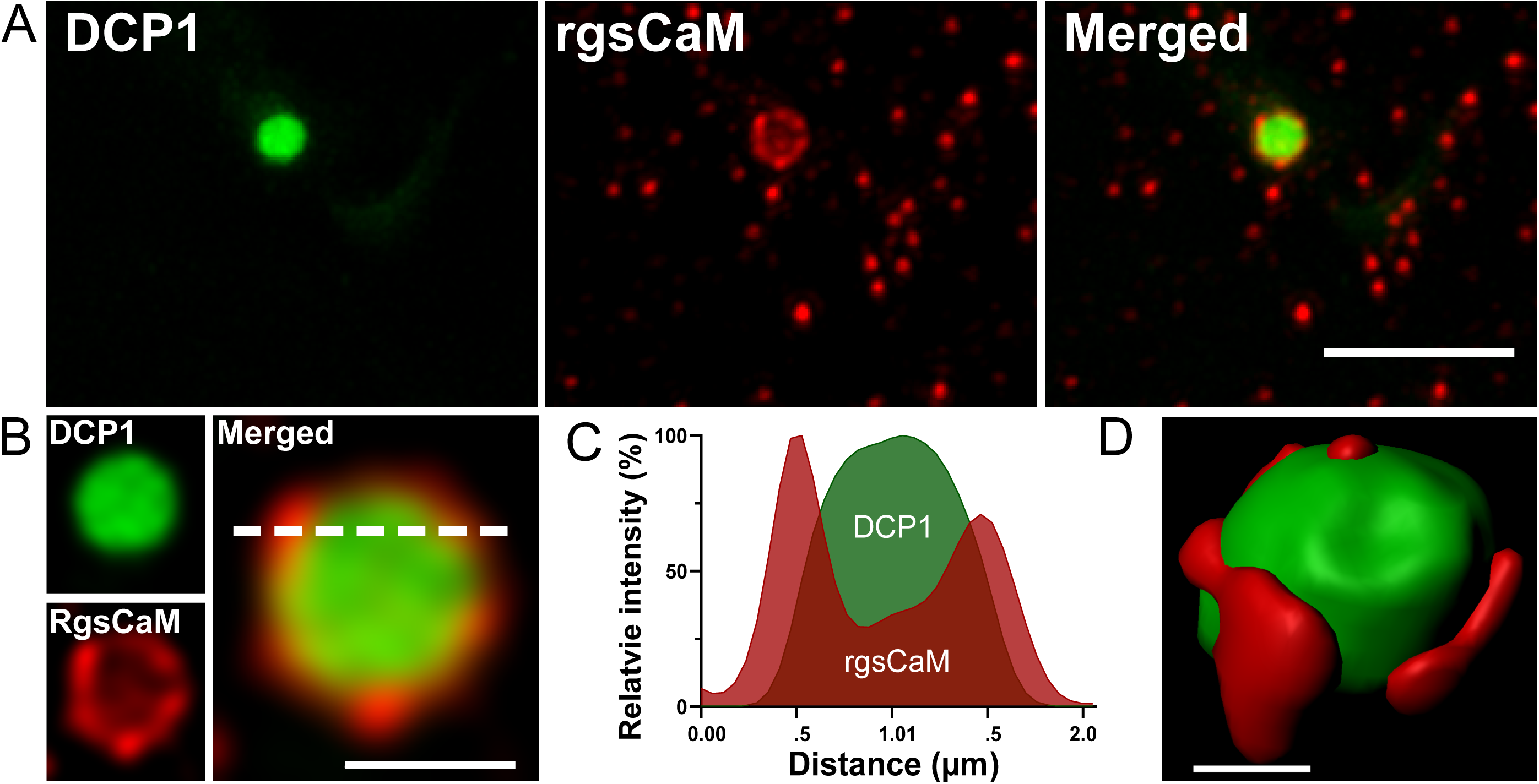
High resolution model of rgsCaM and DCP1 co-localization. **A)** A higher resolution two-dimensional projection from 27 optical sections (0.3 µm step size) of a representative DCP1 processing body structure from a *N. benthamiana* cell co-transfected with *DCP1-CFP* and *rgsCaM-YFP* as described in Fig. 4. Leaf sections were subjected to submergence-induced hypoxia before imaging. The image was subjected to standard deconvolution and a smoothing filter was applied in Leica LasX software. Scale bars: top panel is 5 μm, bottom panel is 1 μm. **B)** A line profile for the fluorescence intensities through the dashed line shown in panel A. **C)** 3D model constructed from the dataset shown in A using the Imaris Software (see also Fig. S2). Scale bar is 0.5 µm.

PBs are highly dynamic structures that interact and dock with other mRNP structures (e.g., SG) within the cytosol, an event that is proposed to facilitate the transfer of mRNA and other components between these two structures (Protter and Parker, 2016). The dynamic nature of the association of rgsCaM bodies with PBs is apparent from the results of live-cell confocal microscopy that show rgsCaM-YFP granules move to, dock, and stably forming a physical association with DCP1 PBs (Fig. S3). These findings suggest that rgsCaM bodies traffic to SGs and PBs and become bound at the surface, accumulating during hypoxia.

### SGS3 localizes to hypoxia-induced cytosolic foci and colocalizes with rgsCaM

Previous observations show that rgsCaM associates with the protein product of *Suppressor of Gene Silencing* 3 (Li et al., 2017). SGS3 associates with cytosolic granule structures referred to as SGS3/RDR6 bodies (Kumakura et al., 2009), or siRNA bodies (Jouannet et al., 2012). To investigate further the localization of SGS3, and the relationship of these bodies to rgsCaM structures, SGS3 was expressed as a CFP fusion in *N. benthamiana*. While a small number of SGS3 granules were present at 10 min after slide preparation, the SGS fluorescence signal was largely diffuse (Fig. 6). Upon submergence, the SGS3 signal shows a time dependent redistribution to well-defined cytosolic foci (Fig. 6). These observations show that SGS3 granule-like structures, similar to SG and rgsCaM bodies, are induced by submergence and hypoxia.

**Fig 6.**
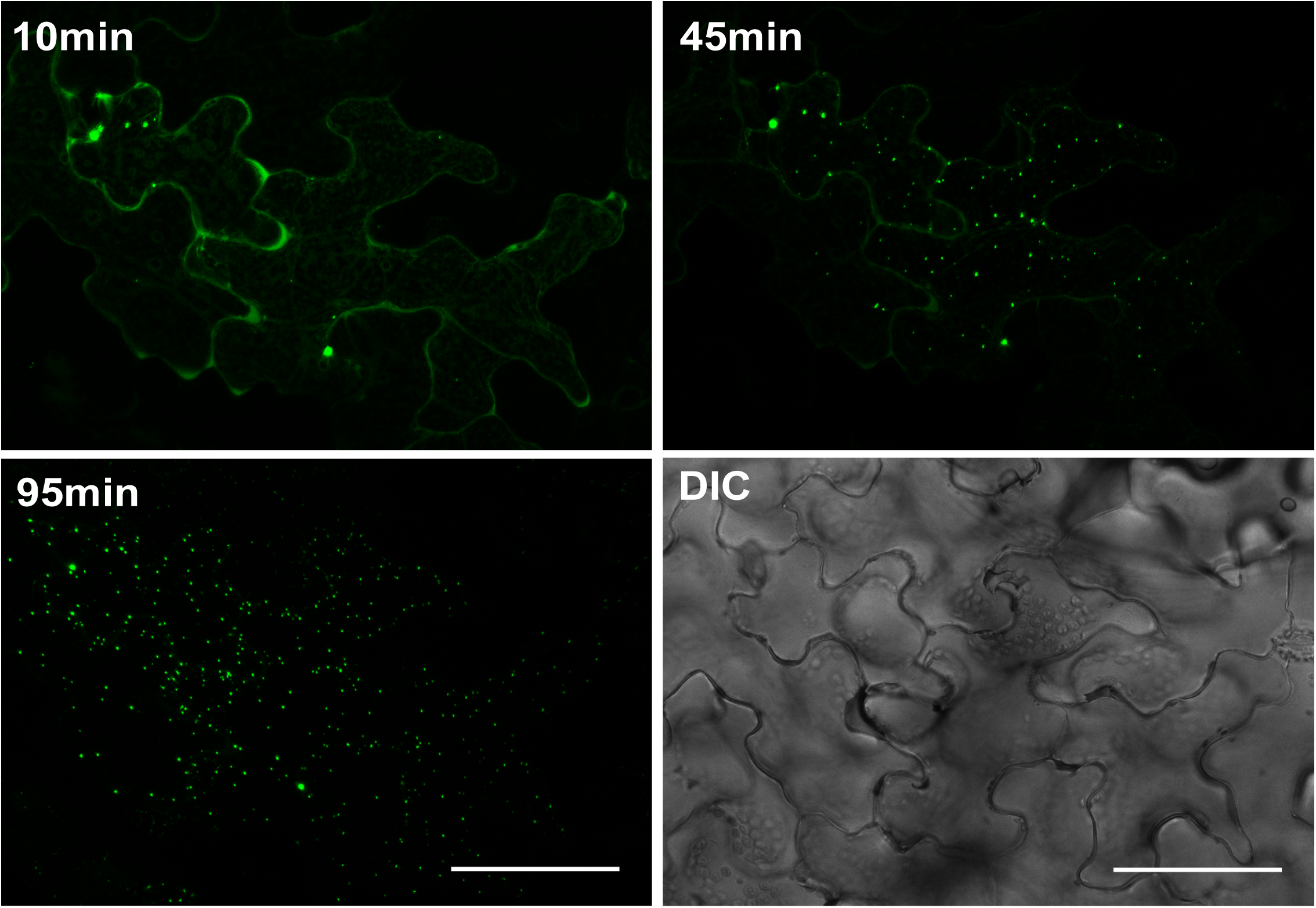
Time course of formation of SGS3-CFP cytosolic foci in response to submergence-induced hypoxia. *N. benthamiana* leaves were transfected with a *SGS3-CFP* construct by *Agrobacterium* infiltration and was imaged over the indicated times after submergence. Scale bar is 50 µm.

To investigate whether rgsCaM and SGS3 co-localize within these structures, co-transfection experiments were performed (Fig. 7). SGS3 bodies are readily observed earlier in the hypoxia time course whereas rgsCaM fluorescence remains diffuse (Fig. 7, top panel). However, over time, rgsCaM redistributes to SGS3 granule structures (Fig. 7, bottom panel) with high Manders co-efficients of co-localization (MCC_SGS3:rgsCaM_ = 0.87, while the MCC_rgsCaM:SGS3_ = 0.95). High resolution imaging with the Leica Lightning de-convolution system revealed the rgsCaM signal is virtually superimposable with the SGS3 signal, with a slightly higher rgsCaM intensity calculated at the periphery of the granule (Fig. 8B-C). Taken together, the data show that co-expression of rgsCaM with SGS3 results in a time-dependent co-localization of the two signals to granule-like structures.

**Fig 7.**
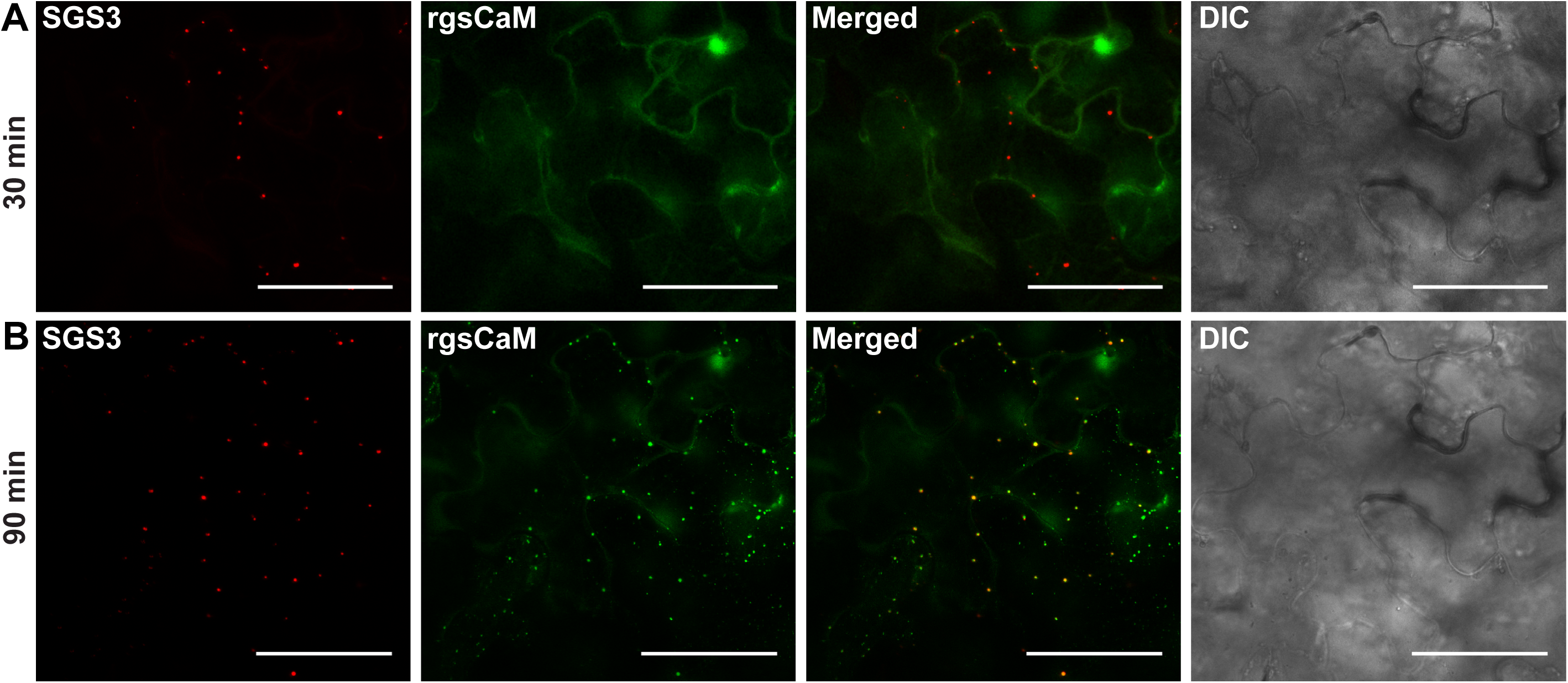
SGS3 and rgsCaM co-localize into cytosolic granule-like structures during hypoxia with different kinetics. *N. benthamiana* leaves were co-transfected with constructs that express rgsCaM-YFP and the stress granule marker SGS3-CFP and leaf sections were subjected to hypoxia as in Fig. 6. Images are 2D projections of 6 optical sections with a step size of 2μm collected at 30 min and 90 min after submergence. Scale bars are 50μm.

**Fig 8.**
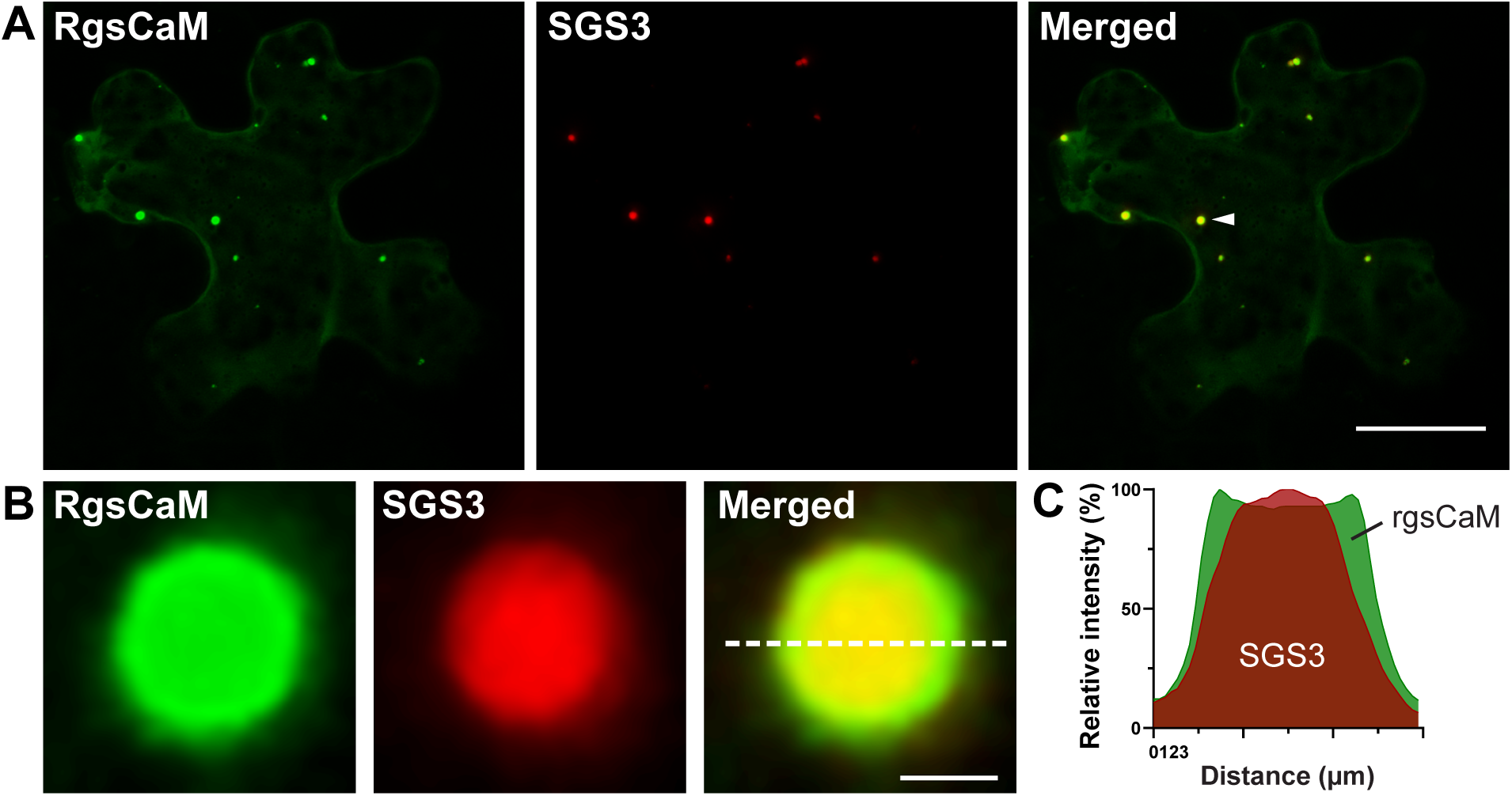
Co-localization of rgsCaM-CFP and SGS3-YFP. **A)** Leaves of *N. benthamiana* were co-transfected with constructs that express rgsCaM-YFP and SGS3-CFP, and leaf sections were subjected to submergence-induced hypoxia. Scale bar is 30 µm. **B)** Enlargement of the structure in A marked with the arrowhead. Images were subjected to a smoothing filter in LAS X and deconvolution was performed with the Leica Lightning system. Scale bar is 1 µm. **C)** Line plot profile of fluorescence intensities through the dashed line in B.

### SGS3 co-localizes with SG

To investigate the potential relationship of SGS3 bodies that form during hypoxia and PB and SG, co-localization experiments were performed with DCP1 and RBP47B markers. Co-transfection of SGS3 with DCP1 results in the formation of cytosolic granule structures that are distinct (Fig. S4) and show low co-localization correlation coefficients (MCC_DCP1:SGS3_ = 0.11 and MCC_SGS3:DCP1_ = 0.12). This is consistent with previous reports that showed that SGS3 bodies are distinct from processing bodies (Kumakura et al., 2009; Jouannet et al., 2012). In contrast, co-transfection of SGS3 with the SG marker RBP47B shows strong co-localization (Fig. 9 A-C) with Manders co-localization co-efficients close to complete superimposition (MCC_RBP47B:SGS3_ = 0.94 and MCC_SGS3:RBP47B_ = 0.88). This suggests that SGS3 bodies and hypoxia-induced SG aggregates may be the same structures. This is supported by previous observations that heat stress also induces the strong relocalization of AGO7 and SGS3 to cytosolic granule structures (Jouannet et al., 2012).

**Fig 9.**
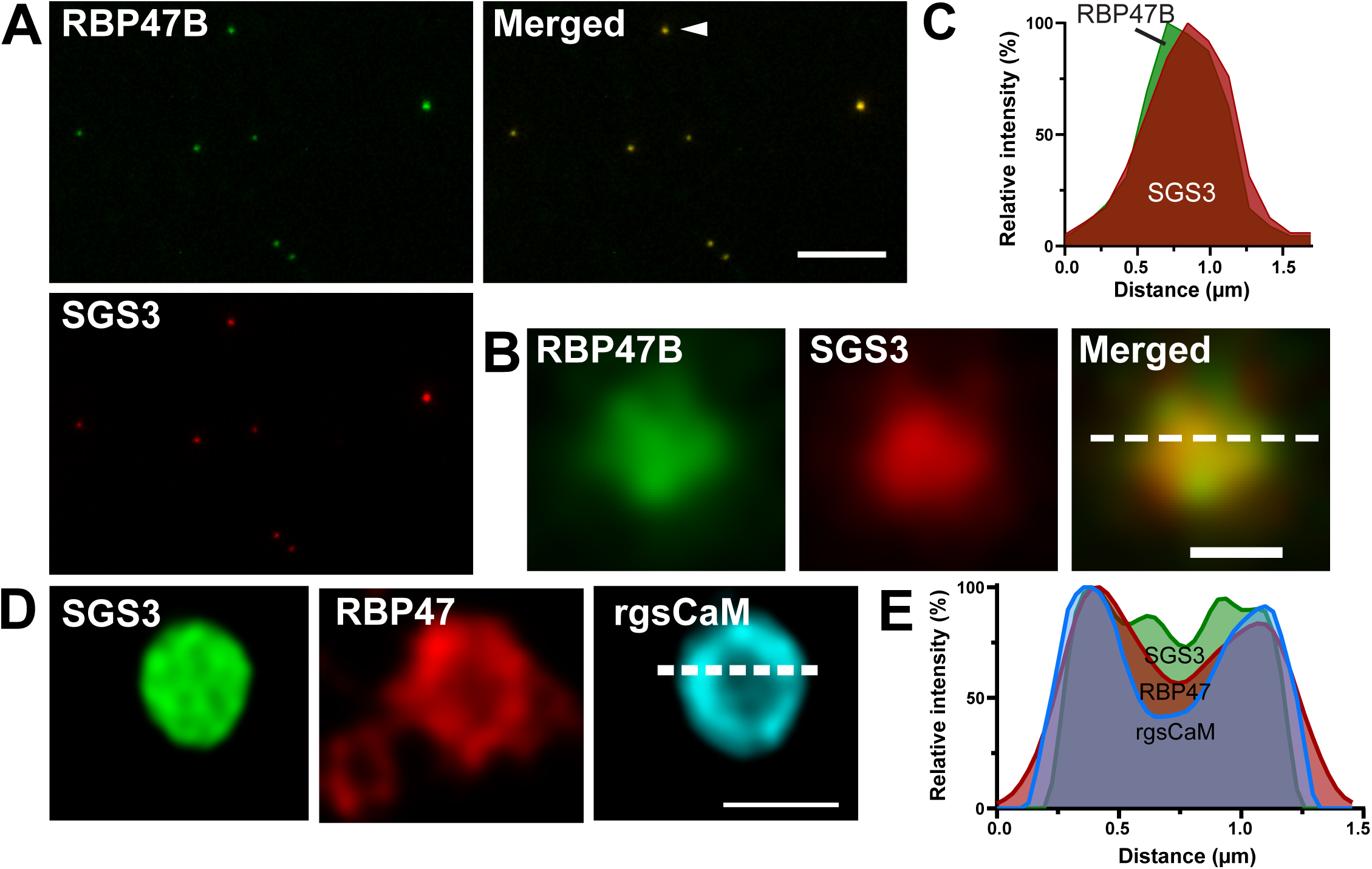
SGS3 and Stress Granule co-localization in *N. benthamiana* cells. **A)** Leaves of *N. benthamiana* were co-transfected with constructs that express SGS3-YFP and the stress granule marker RBP47B-CFP and leaf sections were subjected to hypoxia. Scale bar is 10 µm. **B)** Magnification of the granule marked by the arrowhead in A, with a smoothing filter applied in Leica LAS X. Scale bar is 0.5 µm. **C)** Line plot profile though the dashed line in B. **D)** Triple localization images from *N. benthamiana* leaf epidermal cells co-transfected with SGS3-CFP, RBP47B-RFP and rgsCaM-YFP. Images were taken 5 hr after slide preparation and are 2D projection of 4 optical sections with a Z-step size of 0.13 μm. Scale bar is 1 μm. The Lightning deconvolution system was used and an additional smoothing filter was applied in LAS X. **E)** Line plot through the dashed line in D.

A comparison of the rgsCaM co-localization analyses of SGS3 and RBP47B (Fig. 3 and Fig. 8) reveals that while rgsCaM shows strong correlation with SGS3 bodies, co-localization within the RBP47B SG marker is infrequent with a low level of rgsCaM granule-like structures bound peripherally to SGs (Fig. 3). Based on the previous finding that SGS3 is a direct interaction target for rgsCaM (Li et al., 2017), and the observation here that rgsCaM appears to be recruited to SGS3 bodies over time (Fig. 7), it was hypothesized that overexpression of SGS3 promotes the association of rgsCaM with SGs structures. To test this hypothesis, a triple localization experiment was performed in which *N. benthamiana cells* were co-transfected with expression constructs for SGS3-CFP along with rgsCaM-YFP and RBP47B-RFP. The results show strong co-localization of all three fluorescent signals within RBP47B SGs (Fig. 9D,E). These observations indicate that overexpression of SGS3 causes rgsCaM to localize strongly to SGs.

### RgsCaM interacts with the Autophagy Receptor protein ATG8e

Previous studies have shown that rgsCaM-dependent turnover of binding targets such as SGS3 (Li et al., 2017) and viral suppressors of gene silencing (Nakahara et al., 2012) is mediated by autophagy. To investigate whether rgsCaM co-localizes to autophagosomes, colocalization experiments were performed with the autophagy-related gene product 8e (ATG8e) (Fig. 10). ATG8, similar to its mammalian counterpart LC3, is an ubiquitin-like protein that is conjugated to the amino groups of phosphatidyl ethanolamine during the biogenesis of the autophagosome. ATG8 also serves as an adaptor protein that recruits autophagic cargo to the phagophore during selective autophagy (Stolz et al., 2014; Kellner et al., 2017). As such, fluorescent protein fusions of ATG8e have served as an excellent marker for autophagosomes in plants (Contento et al., 2005). Co-expression of YFP-ATG8e and rgsCaM-CFP in *N. benthamiana* resulted in strong co-localization of the two proteins (Fig. 10A) with high calculated Manders co-efficients (MCCATG8e:rgsCaM = 0.83 and MCCrgsCaM:ATG8e = 0.84).

**Fig 10.**
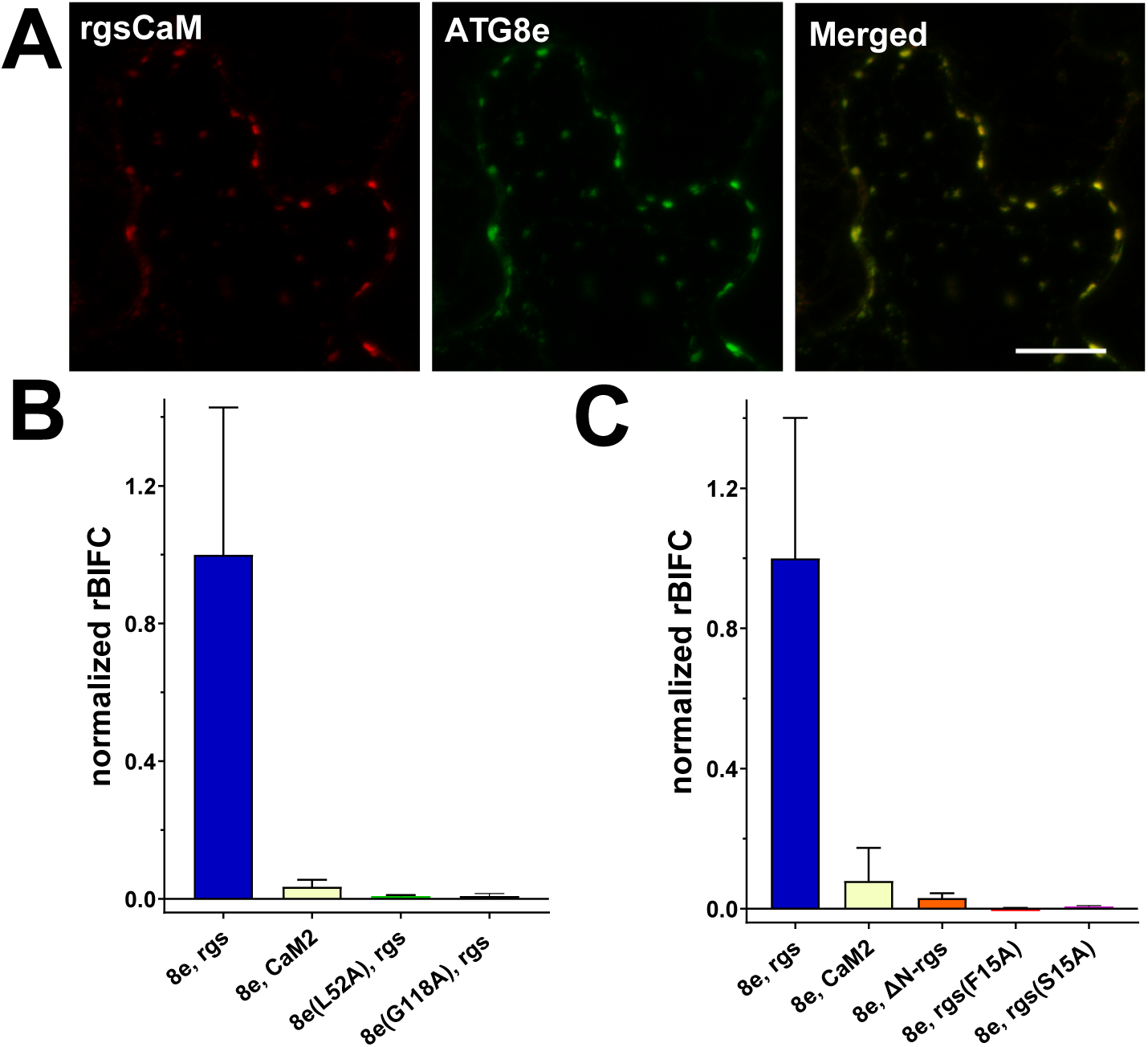
Co-localization and bimolecular fluorescence complementation analysis rgsCaM and ATG8e. **A)** Coexpression of rgsCaM-CFP and YFP-ATG8e in *N. benthamiana* leaves. Images were taken from leaf sections after submergence-induced hypoxia. Scale bars is 20 µm. **B)** Ratiometric BiFC quantification from *N. benthamiana* cells transfected with pBiFC constructs expressing the following protein pairs (see Materials and Methods for details): ATG8e, rgsCaM; ATG8e, AtCaM2; ATG8e, ΔN-rgsCaM; ATG8e, rgsCaM F15A; ATG8e, rgsCaM S16A. **C)** Ratiometric BiFC quantification from *N. benthamiana* cells transfected with pBiFC constructs done as in panel B, except with the following protein pairs: ATG8e, rgsCaM; ATG8e, AtCaM2; ATG8e L52A, rgsCaM; ATG8e G118A, rgsCaM. In both B and C, BiFC signals were standardized as a ratio to constitutive mRFP1 signals and were normalized to the highest value, which was the rgsCaM, ATG8e BiFC pair. The data represent the values from four biological replicates (20 to 30 cells). Error bars show standard deviation. Different letters over histograms represent statistically significant differences (p values <0.0001) based on One-way ANOVA analysis with multiple comparisons.

ATG8 proteins recruit cargo destined for selective autophagy by interaction with an “ATG8 interacting motif” (AIM) consisting of a four amino acid consensus motif [aromatic FWY]-Xaa-Xaa-[aliphatic LIV] on cargo receptor proteins (Birgisdottir et al., 2013; Kellner et al., 2017). Examination of the rgsCaM amino acid sequence revealed a potential AIM motif (residues 15-19: FSRL) within the amino-terminal extension domain that precedes the first EF hand calcium binding site (Fig. S5). To investigate whether rgsCaM and ATG8 physically interact, ratiometric BiFC (rBIFC) was conducted with a single plasmid construct (pBiFC NC *ATG8e*, *rgsCaM)* containing ATG8e as an amino terminal translational fusion with the N-terminal fragment of YFP and rgsCaM as a carboxyl-terminal translational fusion with the C-terminal fragment of YFP. The pBiFC plasmids also encode a constitutive fluorescent reporter protein RFP1 that allows identification of transfected cells, as well as BiFC quantitation and standardization by computing the ratio of the BiFC and RFP1 signals (Grefen and Blatt, 2012).

Transfection of *N. benthamiana* with pBiFC NC *ATG8e*, *rgsCaM* showed a robust BiFC signal (Fig 10 and Fig S6) that suggests that the two proteins interact in planta. As a negative control to test the specificity of this interaction, BiFC experiments were conducted in which rgsCaM was replaced with Arabidopsis calmodulin 2 (CaM2). CaM2 is a conventional calmodulin (i.e., not in the rgsCaM-like clade). When expressed in *N. benthamiana*, CaM2 accumulates as a diffuse cytosolic signal that is distinct from cytosolic granules (Fig. S7). Transfection of *N. benthamiana* with pBiFC NC *ATG8e*, *CaM2* resulted in the near complete absence of the BiFC signal with the signal intensity less than 10% of the rgsCaM-ATG8e pair (Fig. 10).

To investigate further whether rgsCaM has the properties of an AIM cargo receptor, a series of site-directed mutant versions of ATG8e were examined. To disrupt the ability of ATG8e to interact with the AIM domain of proteins, a substitution of leucine 52 with alanine was generated. This leucine residue is conserved in LC3 and ATG8 proteins, and forms part of the W hydrophobic site that interacts with the first aromatic residue in the AIM motif (Noda et al., 2010). The L52A substitution resulted in the loss of the BiFC signal suggesting that an intact AIM interacting site in ATG8e is necessary for rgsCaM interaction (Fig. 10B). Another ATG8e mutant (G118A) mutation also resulted in the loss of the BiFC signal (Fig. 10B). This substitution at the C-terminus abrogates ATG8e lipidation, a process that is critical for localization of the protein to the growing phagophore membrane (Stolz et al., 2014; Marshall and Vierstra, 2018). Overall, the data suggest that the integrity of the AIM interaction site and ATG8 lipidation are essential for rgsCaM interaction.

To test whether the putative AIM motif within the amino terminal domain of rgsCaM is necessary for its interaction with ATG8, rBiFC experiments were conducted with an rgsCaM N-terminal truncation. ΔN-rgsCaM was engineered to remove the N-terminal amino acids 1-42 without disruption of the EF hands within the calcium-sensor domain (Fig. S5). ΔN-rgsCaM fails to produce an rBIFC signal (Fig. 10B) supporting the requirement of the N-terminal extension of rgsCaM for interaction with ATG8. To narrow down the region necessary for interaction, two site-directed mutants of rgsCaM within the putative AIM domain were tested. The F15A mutant has a substitution of a phenylalanine to alanine in the W region of the putative AIM domain, whereas the S16A mutant has a substitution of an alanine for a serine at a position adjacent to the aromatic residue that is conserved in all rgsCaM sequences within the Solanaceae (Fig. S5). Both substitutions result in loss of interaction with ATG8e (Fig 10B) indicating that the integrity of the putative AIM domain is necessary for ATG8 interaction. Analysis of the subcellular localization of the two AIM domain mutants (Fig. S8) show that they are identical to wild type rgsCaM in their ability to localize to hypoxia-induced granules. This suggests that disruption of the N-terminal AIM domain selectively affects association with ATG8 but not assembly into hypoxia-induced bodies.

The data suggests that the amino-terminus of rgsCaM contains a bona fide AIM domain that is characteristic of selective autophagy cargo receptors, and which is necessary for binding to ATG8e. In light of previous findings that rgsCaM drives the autophagic turnover of SGS3 (Li et al., 2017) and viral proteins (Nakahara et al., 2012), and the time-dependent association of rgsCaM with the periphery of RNA granules (Fig. 3 and 4), rgsCaM may target mRNP particles for autophagic turnover in a process similar to granulophagy (Protter and Parker, 2016). The association of mRNPs and rgsCaM with autophagosomes was investigated further by triple localization experiments utilizing high-resolution imaging with super-resolution (Leica Lightning) deconvolution. DCP1-CFP PBs co-localize with rgsCaM-mScarlet and YFP-ATG8e in a putative autophagosome (Fig. 11). Spatial resolution of the fluorescent signal shows co-localization of ATG8e and rgsCaM with highest intensity at the PB periphery (Fig. 11).

**Fig. 11.**
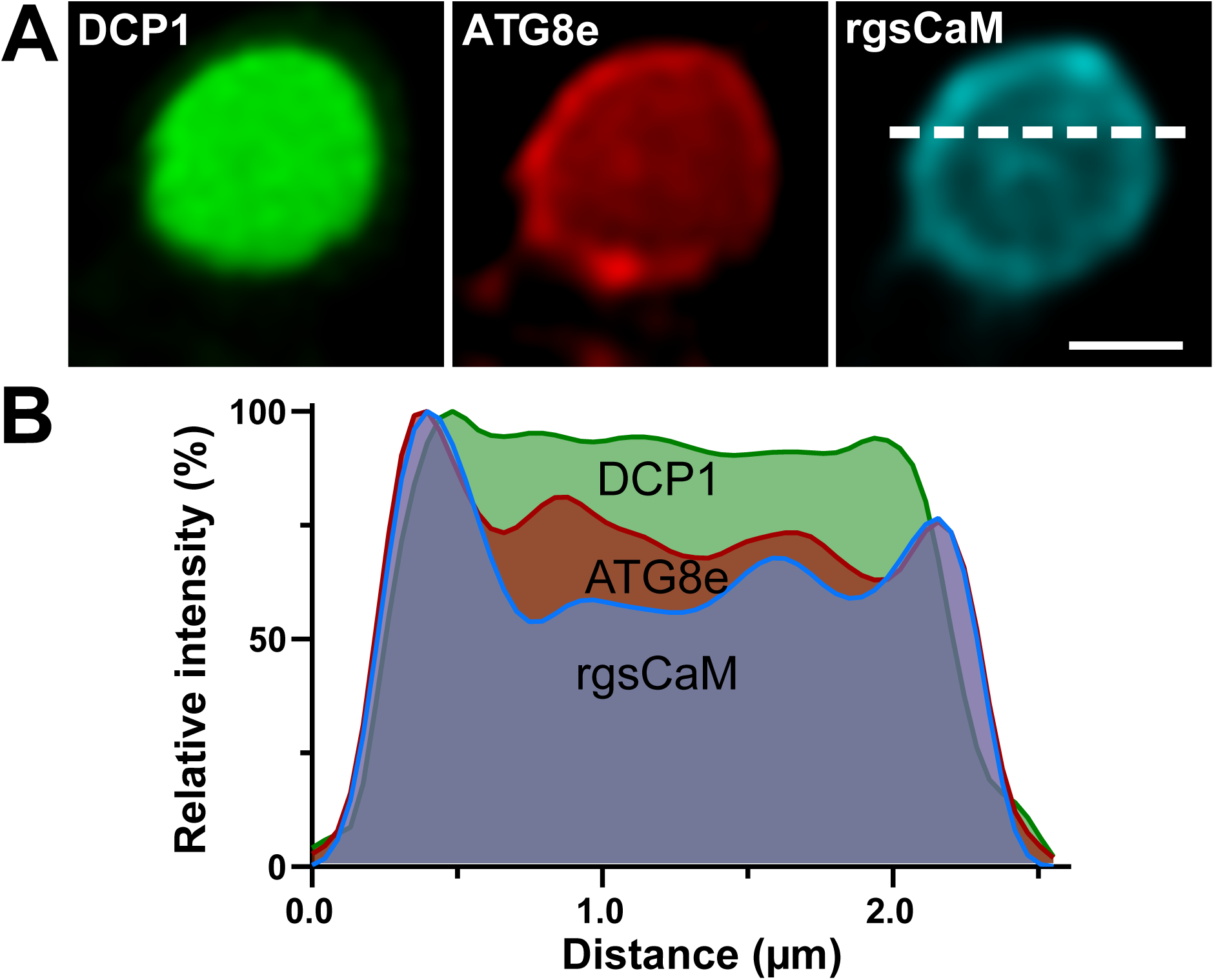
Triple co-localization analysis of rgsCaM, ATG8e, and the PB marker DCP1 in hypoxic *N. benthamiana* leaves. **A)** Co-localization of DCP1-CFP, YFP-ATG8e and rgsCaM-mScarlet transfected *N. benthamiana* leaves after submergence-induced hypoxia. Images are a 2D projection of 9 optical sections with a Z-step size of 0.2 μm. The scale bar is 1 μm. These images were generated using the Lightning deconvolution system. The smooth rendering filter was applied in LAS X. **B)** Line plot through the dashed line in panel A generated from the unfiltered data.

## Discussion

### RgsCaM localizes to independent hypoxia-induced cytosolic bodies that interact with SG and PB

In response to cellular stresses and changing environmental conditions, eukaryotic organisms reprogram their protein translational machinery and mRNA resources. Within the cytosolic network, this occurs in part through the “RNA cycle” through which mRNA is partitioned between translating (polysomes) and non-translating (SG and PB) ribonucleoprotein particles (Protter and Parker, 2016; Chantarachot and Bailey-Serres, 2018; Ivanov et al., 2019). SGs are non-membranous cytosolic organelles that form under conditions in which mRNA translation is impaired or restricted. SGs consist of mRNA bound in an arrested pre-initiation (48S) complex comprised of small ribosomal subunits, translation initiation factors, and a myriad array of RNA binding proteins and other regulatory factors (Kedersha et al., 2013; Buchan, 2014; Protter and Parker, 2016). SGs are dynamic, and interact with and exchange mRNA and protein components with other mRNP complexes including PBs, cytosolic structures that contain machinery for mRNA degradation and posttranscriptional gene silencing (Kedersha et al., 2005; Buchan, 2014). SG are proposed to serve as “mRNA triage” centers that sort mRNA for various cellular fates based on cell needs and environmental cues (Anderson and Kedersha, 2008).

In plants, submergence/hypoxia stress leads to the rapid decline in translating mRNA polysomes (Branco-Price et al., 2008), and the concomitant accumulation of cytosolic mRNP SG granules (Sorenson and Bailey-Serres, 2014). Among the proteins that accumulate within Arabidopsis SG during low oxygen stress is the rgsCaM-like protein CML38 (Lokdarshi et al., 2016). In the present work, it is shown that rgsCaM also relocalizes to cytosolic granule-like bodies in response to submergence stress. However, unlike CML38, rgsCaM bodies localize to structures that are smaller and distinct from SG and PB, but which localize to and dock to the surface of these structures. The identity of these rgsCaM bodies and their relationship to larger SG and PB remains to be determined. As noted below, co-expression of rgsCaM with the rgsCaM-binding protein SGS3 results in integration into the SG structure. The docking of mobile rgsCaM bodies with SGs and PBs may represent a precursor to this structure. During the assembly of SG, smaller, more mobile sub-micrometric granules are trafficked on the microtubule cytoskeleton to larger less mobile SGs for incorporation (Chernov et al., 2009; Loschi et al., 2009; Gutierrez-Beltran et al., 2015; Jain et al., 2016).

### Suppressor of Gene Silencing3 localizes to SG and induces rgsCaM association with these structures

The rgsCaM-interacting protein SGS3 also showed time-dependent localization to cytosolic foci upon submergence. Previous work has demonstrated the localization of SGS3 to similar structures designated “SGS3/RDR6 bodies”(Kumakura et al., 2009) or “siRNA bodies” (Jouannet et al., 2012) that contain SGS3, RDR6, and AGO7. The potential relationship of siRNA bodies to SG was noted by (Jouannet et al., 2012) who observed that heat stress induced the strong accumulation of AGO7 and SGS3 with the SG protein UBP1 within cytosolic granules. In the present work, it is shown that hypoxia induces the re-localization of SGS3 from a diffuse cytosolic localization to cytosolic granules that are virtually superimposable with the SG marker RBP47B. This suggests that SGS3 bodies and SGs may represent the same or closely related particles.

Co-expression of rgsCaM with SGS3 also alters the localization pattern of rgsCaM. Whereas rgsCaM is largely localized to distinct and independent cytosolic structures, co-expression with SGS3 results in a re-localization and integration of rgsCaM within SGs. The SG localization of rgsCaM occurs with different kinetics, with SGS3 granules forming before recruitment of rgsCaM. Previous studies of SG show that these structures show a complex architecture with “core” and “shell” components with different functions and stabilities within the granule (Jain et al., 2016; Protter and Parker, 2016). Examination of the fluorescent signals within SGs from the triple localization experiment (Fig. 9 D and E) show a distinct distribution of the fluorescence signals of SGS3, and rgsCaM, with SGS3 appearing to be predominantly associated with the core of the structure and rgsCaM showing a greater distribution to the periphery (presumably the “shell”). This localization and time-dependent recruitment of rgsCaM to SGS3 granules supports the proposal that SGS3 binds and assists in the recruitment of rgsCaM to SG. The signal that triggers re-localization of rgsCaM to SGS3 granules is not known. However, since previous work shows that the calcium sensor domain containing EF hands I and II is necessary for rgsCaM interaction with SGS3 (Li et al., 2017), a calcium signal is a logical candidate.

### RgsCaM associates with the phagophore protein ATG8 through an AIM domain

What is the significance of rgsCaM association with SGS3 and granule structures? Potential insight comes from the work of (Li et al., 2014; Li et al., 2017) and the role of rgsCaM as a target of the geminivirus suppression of host PTGS. Infection of *N. benthamiana* with geminivirus induces the expression of rgsCaM which mediates the suppression of host RNA silencing by repressing the expression of RDR6 (Li et al., 2014). Subsequent studies revealed that rgsCaM mediates RDR6 turnover by binding to the SGS3 protein that forms a complex with RDR6 (Li et al., 2017). RgsCaM-mediated turnover of SGS3 and RDR6 is suppressed by autophagy inhibitors, and it was proposed that rgsCaM interaction with SGS3 triggers autophagic turnover of SGS3:RDR6 (Li et al., 2017).

Autophagy is a mechanism by which cells degrade and recycle material from the cytosol in response to developmental cues, stresses (unfolded protein response, oxidative stress, heat stress, salt/osmotic stress, hypoxia), and nutrient availability (Marshall and Vierstra, 2018). Autophagy involves a specialized membranous structure called a phagophore, which expands and fuses around its cargo, enveloping it within a sealed double membrane autophagosome vesicle. Autophagosome biogenesis is a complex process involving the coordinated interaction of an array of “autophagy related proteins” (ATGs). Among these is ATG8, a ubiquitin-like protein, that plays a central role in selective autophagy by recruiting specific cargo to the growing phagophore (Michaeli et al., 2016; Johansen and Lamark, 2019). During autophagosome biogenesis, ATG8 is cleaved near its C-terminus by ATG4 peptidase to expose a C-terminal glycine that leads to covalent conjugation (“lipidation”) to phosphatidyl ethanolamine within the phagophore membrane (Stolz et al., 2014; Michaeli et al., 2016; Marshall and Vierstra, 2018). ATG8 then recruits cargo to the growing phagophore through “cargo adaptor proteins” that contain an ATG8-interacting motif (AIM) (Johansen and Lamark, 2019).

The present study presents evidence that rgsCaM contains an AIM domain within its amino terminal domain that is conserved within rgsCaM-like proteins in the Solanaceae family (Fig. S5). Ratiometric BiFC shows that ATG8e and rgsCaM physically interact, and that mutations within the AIM domain abolished this interaction. Conversely, ATG8e substitutions that disrupt the interaction site for AIM domain interaction or its ability to be lipidated, also eliminate ATG8e:rgsCaM interaction.

### RgsCaM and related proteins as targets for autophagy during viral infection and abiotic stress

Taken together with previous evidence that rgsCaM mediates the autophagic turnover of SGS3 and RDR6 (Li et al., 2017), and the time dependent recruitment of rgsCaM to SGS3 granules observed here, it is proposed that rgsCaM may serve as a cargo adaptor protein that mediates the turnover over of SGS3-associated mRNP granules. This process would be similar to “granulophagy”, a selective autophagy process that the leads to the degradation of SG, PB, and other RNA granules (Buchan et al., 2013; Frankel et al., 2017). Granulophagy was first discovered in yeast, but also is widely observed in metazoan cells, and defects in this process has been linked to disease states such as Amyotrophic Lateral Sclerosis (Buchan et al., 2013; Krisenko et al., 2015; Monahan et al., 2016; Chitiprolu et al., 2018). While the autophagic degradation of RNA granules in plants is less well studied, granulophagy has been proposed as part of the plant host’s defense response during viral infection (Hafren et al., 2018), and in general has been postulated to be a component of plant RNA quality control (Yoon and Chung, 2019).

A potential mechanism for rgsCaM and related proteins in the regulation of autophagy and RNA in response to viral infection and abiotic stresses is shown in Fig. 12. With respect to abiotic stress, the expression of rgsCaM-like proteins are induced by a variety of environmental stresses, including osmotic and salinity stress, wounding, and flooding/hypoxia (Vanderbeld and Snedden, 2007; Xu et al., 2011; Tadamura et al., 2012; Lokdarshi et al., 2016). For example, the Arabidopsis rgsCaM-like protein CML38 is a core-hypoxia response protein that is induced over 300-fold during flooding or hypoxia stress, and which accumulates within SGs (Lokdarshi et al., 2016). SGs are induced during sustained hypoxia (Sorenson and Bailey-Serres, 2014) due to the decrease in translating polysomes (Branco-Price et al., 2008) associated with the energy crisis induced by anaerobic stress. CML38-associated granules are degraded during subsequent return to normoxic conditions (Lokdarshi et al., 2016). Similar to rgsCaM (Fig. 12), CML38 has the same domain organization with two EF hands containing calcium sensor domains with an extended amino-terminal domain that contains a potential AIM site (Fig. S5). Flooding stress induces the expression of autophagy related genes (ATGs), increases the number of autophagosomes, and increases autophagic degradation (Chen et al., 2015). ATG gene mutants show increased sensitivity to hypoxia indicating that autophagy is an important component of hypoxia survival (Chen et al., 2015), and plant proteins with AIM domains are targets for selective autophagy during hypoxia [e.g., S-nitroso-glutathione reductase 1 (Zhan et al., 2018)]. While the work here shows that rgsCaM associates with hypoxia-induced granules in *N. benthamiana*, its remains unknown whether rgsCaM and related proteins are part of the hypoxia adaptive response in the Solanaceae. One observation of interest is that a number of Solanaceae rgsCaM proteins possess amino-terminal cysteine residues (Fig. S5) that are characteristic of proteins regulated by hypoxia through the N-degron pathway (Masson et al., 2019).

**Fig. 12.**
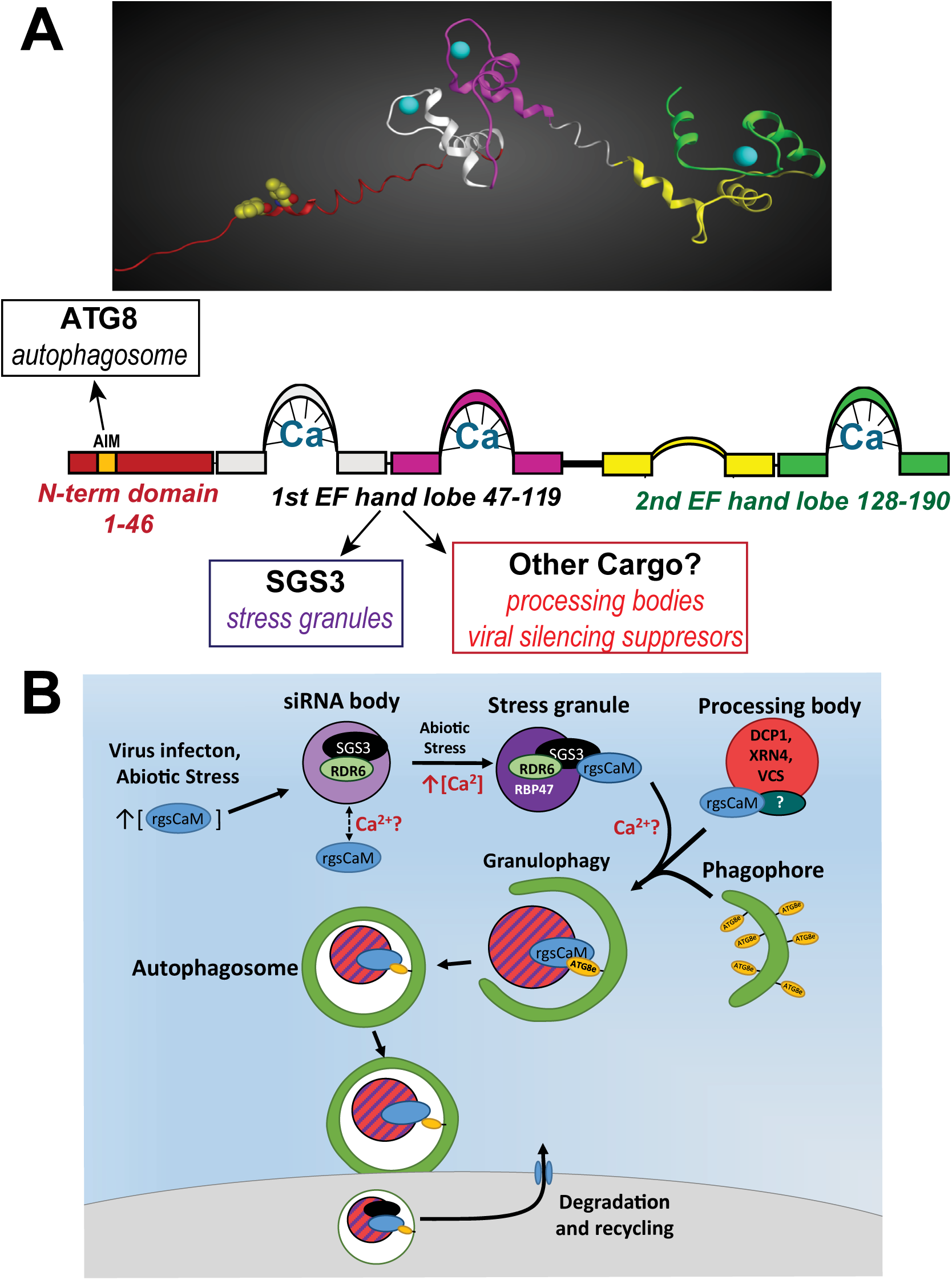
Model for rgsCaM as a regulator of autophagic turnover of SGS3 and other cargo. **A.** *Top panel*, domain model for rgsCaM based on homology modeling with a calcium-calmodulin template (pdb 1cll). The bilobal structure with EF hands I (white) and II (magenta) comprising the 1^st^ calcium binding lobe (residues 47-119) and EF hands III (yellow) and IV (green) comprising the 2^nd^ calcium binding lobe. The N-terminal domain is simply shown as an extended structure with the predicted AIM domain residues shown in space filled representation. *Bottom panel*, diagram showing the predicted sites of binding of rgsCaM to ATG8, as well as to SGS3 and other cargo (PB and viral suppressors of gene silencing). **B.**Model for rgsCaM mediated turnover of SGS3, RNP granules, and other cargo. RgsCaM and rgsCaM-like proteins (e.g., CML38) are induced during viral infection and selective abiotic stresses (e.g., hypoxia). Suppressor of gene silencing 3 (SGS3) resides in siRNA bodies as well as stress granule structures in complex with RNA-dependent RNA-polymerase 6 (RDR6). SGS3 recruits cytosolic rgsCaM to stress granules, and mediates the turnover (and suppression of secondary siRNA silencing) through targeting of SGS3 granules to autophagosomes (granulophagy) by rgsCaM interaction with ATG8. RgsCaM also interacts with processing bodies through interaction with an unknown PB protein target. In addition to degradation of the SGS3/RDR6-mediated siRNA production machinery, autophagic degradation of mRNPs can recycle material during biotic and abiotic stress, and may control mRNA homeostasis and expression.

With respect to viral infection, viral suppressors of silencing (e.g., potyviral HCPro and geminiviral βC1), induce the expression of rgsCaM or rgsCaM-like proteins (Anandalakshmi et al., 2000; Endres et al., 2010; Li et al., 2014; Yong Chung et al., 2014). SGS3 binds to the calcium sensor domain of rgsCaM that contains EF hands I and II [Fig. 12, (Li et al., 2017)]. It is proposed here that the AIM domain in the N-terminal extension of rgsCaM mediates the recruitment of SGS3 “marked” siRNA bodies or SGs to ATG8 within the growing phagophore (Fig. 12). The subsequent degradation of the SGS3/RDR6 complex/granules via autophagy limits the production of secondary viral siRNAs. Notably, SGS3 degradation via autophagy is also induced by other viral protein components (e.g., the viral genome-linked protein VPg from potyviruses) (Cheng and Wang, 2017), indicating multiple pathways through which viruses subvert the secondary siRNA defense response of the host.

In contrast to the proposal that rgsCaM serves as a viral target for the suppression of anti-viral siRNA silencing of the plant host, an opposing view for rgsCaM as an anti-viral defense protein has also been proposed (Nakahara et al., 2012). Nakahara et al. (2012) reported that rgsCaM directly binds to viral suppressor proteins (potyviral HCPro and the cucumber mosaic viral 2b protein), and that it mediates the autophagic degradation of these proteins resulting in the loss of viral suppression of the host silencing defense response. These apparent contradictory activities of rgsCaM could possibly be explained by its proposed role as an autophagy cargo receptor, and the fact that rgsCaM is able to interact with multiple protein cargos (e.g., SGS3 or viral proteins, Fig. 12A) with subsequent autophagy leading to opposing effects. During potyvirus infection, HC-Pro induces and becomes localized to specialized RNA granule structures (potyvirus-induced RNA granules, or PGs) that contain components of both SGs and PBs, as well as viral RNA (Hafren et al., 2015; Hafren et al., 2018). Selective autophagic degradation of HC-Pro, and presumably PGs, through the cargo receptor NBR1 is proposed to be part of the anti-viral defense response (Hafren et al., 2018). Whether rgsCaM represents a separate autophagic pathway or participates in some manner within the NBR1 pathway is unknown. As pointed out by Hafren et al. (2018), viral modulation of the autophagy pathway is complex, with both pro-viral as well as anti-viral component. How rgsCaM distinguishes viral (e.g., HC-Pro and other viral suppressors) and host (e.g., SGS3) targets during viral infection and plant defense merits further investigation.

## Materials and Methods

### Molecular cloning techniques and procedures

Expression constructs with translational fluorescent protein fusions driven by the Cauliflower Mosaic Virus (CaMV) 35S promoter were generated in pEarleyGate vectors: pEG101 (C-terminal YFP), pEG102 (C-terminal CFP), pEG104 (N-terminal YFP), and pK7RWG2 (C-terminal RFP) (Karimi et al., 2005). All are Gateway-compatible destination vectors with *attP1* and *attP2* recombination sites. Donor clones were generated in pDONR207 (Invitrogen).

*Nicotiana tabacum rgsCaM* cDNA (Anandalakshmi et al., 2000) and Arabidopsis *ATG8e* cDNA (Contento et al., 2005) were amplified using gene specific primers that add the attB1 recombination site to the 5’ end and the attB2 site to the 3’ end (Supplemental Table S1). For *SGS3* and *DCP1* cloning, cDNA was generated with the SuperScript III first strand cDNA synthesis system (ThermoFisher) using total RNA from 1-2 week old *Arabidopsis thaliana* Col-0 seedlings as a template followed by amplification with gene specific primers with the *attB1* recombination and the *attB2* recombination sites (Supplemental Table S1). Amplified PCR products were isolated and purified by gel electrophoresis, and were recombined with the Gateway-compatible donor vector pDONR207 in a BP clonase II (Invitrogen) reaction as previously described (Lokdarshi et al., 2016). The desired cloned products were confirmed by DNA sequencing, and donor clones were recombined with one of the localization vectors above in an LR clonase II reaction per manufacturer’s protocol (Invitrogen). The following Gateway-compatible clones were generated and used in this study: pEG 101 *rgsCaM-YFP*, pEG 102 *rgsCaM-CFP*, pEG 102 *DCP1-CFP*, pEG 101 *SGS3-YFP*, pEG 102 *SGS3-CFP*, pEG 104 *YFP-ATG8e*, and pK7RWG2 *RBP47B-RFP*. pEG 102 *RBP47B*-CFP was generated as previously described in (Lokdarshi et al., 2016).

A translational fusion construct of *rgsCaM* with the *mScarlet* fluorescent reporter protein (Bindels et al., 2017) was prepared in pBin19 (Frisch et al., 1995). *Xba*I and *Xma*I sites were introduced into the *Nt-rgsCaM* open reading frame at the 5’ and 3’ ends respectively by PCR using gene specific primers with these sites (Table S1). *Nt-rgsCaM* was then subcloned into these sites within a shuttle plasmid (a gift from Andreas Nebenführ, University of Tennessee – Knoxville) to generate the *mScarlet* translational fusion under the control of the *35SCaMV* promoter. A fragment containing *35S:rgsCaM-mScarlet* was excised by digestion with *Sac*I and *Hind*III, and subcloned into pBin19.

pBiFCt-2in1 plasmids (Grefen and Blatt, 2012) were used for ratiometric BiFC. Donor clones were generated in pDONR 221 plasmids containing either *attP1* and *attP4* sites, or *attP3* and *attP2* sites. BiFC pairs were generated by using Multisite LR clonase II *plus* (Invitrogen) with each reaction containing the desired pDONR221 donor construct pairs and a pBiFC 2-in1 plasmid (3:3:1 molar ratio) (Grefen and Blatt, 2012). Site-directed mutagenesis was conducted on pDONR 221 plasmids containing Arabidopsis *ATG8e* and on pDONR 221 plasmids containing *Nt-rgsCaM*. Mutagenesis was performed according to the “’round the horn” protocol (OpenWetWare). All mutagenesis primers and corresponding mutations are described in Table S1. An N-terminal truncation of rgsCaM that removes amino acids 1-42 was also generated. The coding region of *Nt-rgsCaM* corresponding to amino acids 43-190 was amplified from pDONR207 *Nt-rgsCaM* using an rgsCaM ΔN attB1 forward primer and the rgsCaM attB4 reverse primer (Table S1) and was recombined into pDONR 221 to generate a donor plasmid containing *ΔN-rgsCaM*.

Sanger DNA sequence analysis of all plasmid constructs was performed on a Applied Biosystems 3730 Genetic Analyzer at the University of Tennessee Genomics Core Facility.

### Plant growth, transfection, and stress treatments

*Nicotiana benthamiana* seeds were germinated in Lambert LM-GPS soil and were grown in an environmental growth chamber with long day conditions (16h light at 25°C/8h dark at 23°C) and 50% humidity under a light intensity of 100 µmol m^-2^ s^-1^. Plants that were between four to five weeks old (prior to flowering) were used for transfections. Binary vectors were introduced into *Agrobacterium tumefaciens* strain GV3101 (Koncz and Schell, 1986) by electroporation (Jones, 1995) or heat shock (Hofgen and Willmitzer, 1988), and selected colonies were grown at 28°C for 24-48 hr in LB broth with 50 µg/mL rifampicin, 50 µg/mL gentamycin and either 50 µg/mL kanamycin (for pEarleyGate plasmids) or 100 µg/mL spectinomycin (for pBiFCt-2in1 plasmids). Cells were collected by centrifugation at 3,000 x *g* for 10 min and pellets were resuspended in 10 mM MgCl_2_, 10 mM MES-NaOH, pH 5.7, 200 µM acetosyringone, and infiltration of *N. benthamiana* was performed as in (Brunkard et al., 2015). The inoculum was incubated with slow rotation at 20°C for 2-4 hr to allow acetosyringone-induction of virulence gene expression. For co-infiltrations, cultures containing each plasmid were combined prior to infiltration so that each was equivalent to OD_600_ of 0.4. Combined *A. tumefaciens* cultures were infiltrated into the abaxial side of leaves using a needleless syringe and the infiltrated region was marked. Plants were kept at 20°C overnight on a lab bench, and then transferred to growth chambers under standard long day conditions (16 h light, 22-25°C, 8 h dark, 20-22°C). Imaging experiments were performed two days after infiltration.

Hypoxia stress was administered by submergence of transfected *N. benthamiana* leaf sections as previously described (Lokdarshi et al., 2016). One cm^3^ leaf discs were cut from infiltrated regions adjacent to the site of injection. Tissues were placed in DI water and a brief manual vacuum was administered with a syringe to remove air pockets as described by (Brunkard et al., 2015). Hypoxia was administered by submerging leaf samples in DI water on a slide under a #1.5 coverslip, adaxial face down (Lokdarshi et al., 2016). Imaging was performed on adaxial epidermal cells. Except where indicated in specific figures, submergence stress was administered between 1.5 to 4 hr before imaging.

### Microscopy and imaging techniques

An Leica SP8 laser scanning confocal microscope (Wetzlar, Germany) was used for all confocal microscopy performed in this study. For co-localization experiments, each fluorescence channel was captured sequentially to prevent signal cross-talk. Sequential imaging was performed between scan lines to limit the impact of granule movement. Excitation/emission bandwidth settings for imaging were: 405 nm/425-500nm (DAPI), 470 nm/480-530 nm (CFP), 514 nm/540-575 nm (YFP), 575 nm/600-660 nm (RFP) and

590 nm/600-660 (mScarlet). Chloroplast autofluorescence was captured over 680-740 nm. To reduce chloroplast autofluorescence when imaging CFP, YFP or RFP, time gating was used on the HyD hybrid detectors to exclude emission from the first 0.3 ns following pulsed excitation from the white light laser as described by (Kodama, 2016). Laser strength and gain were adjusted for each image to produce signal just below saturation.

For DAPI staining (4′,6-diamidino-2-phenylindole; Sigma, St. Louis, MO), transfected leaf discs were cut and placed in DI water with 5 µg/ml DAPI for 30 min prior to mounting in DI water for imaging. Co-localization imaging was performed over a Z-series to include the top surface of the cell. Except where indicated in specific figures, the step-size between optical sections was between 1-2 µm. The pinhole size was 0.8-1 AU.

Confocal micrographs were captured in Leica LASX software and uncompressed images were exported and analyzed in ImageJ (Schneider et al., 2012). Brightness was increased to just below saturation, and the built-in smoothing filter in ImageJ was applied to reduce noise. ImageJ was used to add scale bars, false colorations and overlays, and to perform maximum intensity 2D projections. Except where indicated in specific figures, 2D projections shown represent between 20 to 27 optical sections.

Colocalization analysis was performed in ImageJ using the Just Another Colocalization Plugin (JACoP) (Bolte and Cordelieres, 2006). Images corresponding to the two channels were imported and a region of interest was selected that corresponds to the cytosol of an individual cell. JACoP was used to compute Mander’s coefficients (Manders et al., 1992) to obtain quantitative analyses and spatial descriptions of overlapping signals from the two fluorophores. Manders’ coefficients (MCC) are more informative that other co-localization techniques (e.g., Pearson correlation co-efficients) when there are unequal co-localizations, such as one protein strongly co-localizing with the second, while the second also showing independent localization (Bolte and Cordelieres, 2006; Dunn et al., 2011). Two Mander’s coefficients were computed: the first co-efficient quantifies the fraction of the signal from channel A that overlaps with channel B, and the second quantifies the fraction of the channel B signal that overlaps with channel A (Manders et al., 1992).

Data for line plots were obtained by drawing a line through the indicated region in ImageJ and using the built-in plot profile function. The output of brightness values by distance (µm) for each pixel was imported into Excel, converted into percentages based on the highest obtained value by channel, and then imported into Prism 8.1.1 (GraphPad) for figure generation. Standard deconvolution of images was done by using the Huygens Essentials software (Scientific Volume Imaging, Hilversum, The Netherlands) using the standard profile for automatic deconvolution. The Leica Lightning deconvolution system was used in certain experiments (triple localization Fig. 9 and 11, and rgsCaM and SGS3, Fig. 8) to optimize image capture settings for Z-axis step size, resolution and scan speed, and to maximize resolution. The adaptive deconvolution profile setting in the Leica Lightning Deconvolution package was used.

Imaris (Oxford Instruments, Zurich, Switzerland) was used to generate 3D models from deconvolved Z-series signal volumes in selected co-localization experiments. Thresholds were defined for CFP and YFP signals to produce models corresponding to their respective signal volume data. To correct for signal bleed in the Z-axis, the setting for the Z-axis step size in Imaris was adjusted to reflect the spherical shape of the imaged granules (rgsCaM granules were the reference for Fig 3, and the DCP1 granules were the reference in Fig 4).

### Ratiometric bimolecular fluorescence complementation

For BIFC analysis, pBiFC 2-in-1 plasmids (Grefen and Blatt, 2012) expressing the two proteins of interest as nYFP or cYFP fusions were transfected into *N. benthamiana* as described above. The BiFC plasmid also expresses a constitutive mRFP1 that serves as a reference for ratiometric comparison to the YFP BiFC signal. This ratiometric approach allows for quantification of BiFC signals and controls for differences in plasmid copy number among transfected cells. mRFP1 was imaged with excitation/emission bandwidth of 543 nm/560-615 nm, while the eYFP BiFC signal used 514 nm/520-550 nm. Both channels used time gating to exclude the first 0.3 ns to reduce chloroplast autofluorescence (Kodama, 2016). Sequential imaging was performed between scan lines, with a line averaging of four used for each image of a single optical section. Four biological replicates were used, with at least five cells analyzed from each. BiFC clones generated and used in this study include: pBiFC NN *ATG8e*, *rgsCaM*; pBiFC NC ATG8e, *rgsCaM*; pBiFC NN *ATG8e*, CaM2; pBiFC NC *ATG8e*, CaM2; pBiFC NC *ATG8e-L52A*, rgsCaM; pBiFC NC *ATG8e-G118A*, *rgsCaM* and pBiFC NC *ATG8e*, *ΔN-rgsCaM*; pBiFC NC *ATG8e*, *rgsCaM*-*F15A*; pBiFC NC *ATG8e*, *rgsCaM-S16A*, where the two letter code indicates whether the split-YFP is positioned at the N- or C-terminus of each coding sequence. Identical BiFC results were obtained for both N and C terminal fusions of rgsCaM, and only the results with C-terminal fusions of rgsCaM and rgsCaM mutants are shown. A negative control construct (pBiFC NC *ATG8e*, *CaM2*) was generated in which Arabidopsis *calmodulin 2* (*CaM2*) was used in place of *rgsCaM*.

Ratiometric BiFC analysis was similar to (Grefen and Blatt, 2012). In ImageJ, a region of interest was traced to select individual cells. Nuclei were not visible in all images and they were excluded from the ROI when present in case the BiFC complex had different nuclear preference than the mRFP1. ImageJ was used to compute the average brightness of the ROI for YFP and RFP channels, this was then exported to Excel where the eYFP:mRFP1 ratio was calculated for each cell and the values were normalized to the highest signal, which in all experiments was for ATG8e and rgsCaM. Statistical analysis was performed in GraphPad Prism version 8.1.1 (Graph Pad).

### Gene ID and accession numbers

*The* GenBank accession numbers for coding sequences used in this study are: NM_100710.4 (*DCP1*), NM_112800.4 (*RBP47B*), NM_001343817 (*SGS3*), NM_180100.7 (*ATG8e*), NM_180013.3 (*CaM2*), AF329729.1 (*rgsCaM*).

## Acknowledgements

We thank Dr. Andreas Nebenführ and Anna Vick (University of Tennessee, Knoxville) for the kind gift of the pAN990 mScarlet plasmid, and Dr. Tessa Burch-Smith (University of Tennessee, Knoxville) for guidance on transfecting and imaging *N. benthamiana* leaves and for critical comments during the preparation of the manuscript. Microscopy was conducted at the Advanced Microscopy and Imaging Center, a core facility at the University of Tennessee-Knoxville, with support from Dr. John Dunlap and Dr. Andreas Nebenführ. Automated Sanger sequencing was conducted at the Genomics Core facility at the University of Tennessee-Knoxville.

## Supplemental Material

**Fig S1. Fitting of SG-marker RBP47B and rgsCaM imaging data to 3D models.**

**Fig. S2. Fitting of PB-marker DCP1 and rgsCaM imaging data to 3D models.**

**Fig S3. RgsCaM body docking onto DCP1 processing body.**

**Fig S4. SGS3 and DCP1 colocalization in *N. benthamiana* leaves.**

**Fig S5. Domain structures and amino acid sequence of rgsCaM and rgsCaM-like proteins.**

**Fig S6. Bimolecular fluorescence complementation of wt and mutant forms of rgsCaM and ATG8e.**

**Fig S7. Co-localization experiments with Arabidopsis CaM2 and SGS3 in *N. benthamiana* leaves.**

**Fig S8. Localization of rgsCaM AIM mutants in *N. benthamiana*.**

**Table S1: PCR primers for molecular cloning, site-directed mutagenesis and sequencing**

